# The Potential Applications of Site-Directed Mutagenesis for Crop Improvement: A review

**DOI:** 10.1101/2020.10.01.321984

**Authors:** Yilkal Bezie, Tadesse Tilahun, Mulugeta Atnaf, Mengistie Taye

## Abstract

The search for technologies for crop improvement has been a continuous practice to address the food insecurity to the growing human population with an ever decreasing arable land and dynamic climate change around the world. Considering the potential technologies for crop improvement could close the rooms of poverty in developing countries in particular and around the globe at large. This review aimed to assess the site-directed mutation creation methods and to show the potential tools for future crop improvement programs. Site-directed mutagenesis was found to be an efficient process to create targeted mutation on cereal crops, horticultural crops, oilseed crops, and others. Agronomic traits such as yield, quality, and stress tolerance have been improved using site-directed mutagenesis. Besides, selectable marker elimination was also reported from transgenic crops by targeted mutation. Most of the reports on site-directed mutagenesis is focusing on cereal crops (58.339%) followed by horticultural crops (22.92%). Among the four mutagenic tools that have been reported, the CRISPR/Ca9 technology was found to be frequently used (66.67%) followed by TALENs. This tool is potential since it is efficient in creating targeted mutagenesis and less likely off-target effect, so it is repeatedly used in different research works. TALENs were used usually to knockout genes with bad traits. Moreover, the mutation created by mutagenic tools found to be efficient, and the mutated traits proved as it was heritable to generations. Hence, site-directed mutagenesis by the CRISPR/Cas9 system is advisable for agricultural development thereby ensuring food sustainability around the world.

## INTRODUCTION

Agricultural development has always been on the move towards increasing crop productivity. Sustainable use of natural resources must be wisely managed in combination with the enrichment in the knowledge gained from science and technology (1). Global food security continues to be the first issue and plant breeders are obliged to sustain food production to meet the demand of the ever-growing human population around the world (2).

The process of crop improvement has been a fundamental issue for thousands of years ago (3). The ultimate reason to crop improvement is to respond to the huge demand for food for the alarmingly growing human population around the globe (4). Moreover, the ever-greater need for a balanced and healthy diet, there must be an ongoing need to develop improved crops using divers technologies (5, 6). Multifaceted and integrated global strategies are required to ensure sustainable food security through crop improvement programs (3, 7, 8).

Site-directed mutagenesis is one of the recent tools amongst molecular crop improvement technologies (9). The major aim of mutation assisted breeding is to develop and improve well-adapted plant varieties by modifying one or two major traits (10). The development of targeted mutation became a source of genetic variation, in turn, become a resource for plant breeders (11). Therefore, mutation supported plant breeding could play a crucial role in addressing the uncertainties of global climate change and food insecurity challenges (1). Site-directed mutagenesis is aimed at precise change of any coding sequence in *vitro or in vivo*. Site-directed mutagenesis could be produced using different methods. *In vitro* targeted mutation could be created by gene vector-based method or PCR based method (12, 13).

Another method of the site-directed mutation creation method is gene editing using programmable site-directed nucleases (SDN), which are promising for new plant breeding techniques. This method could be achieved by generating a small deletion or insertion at a precisely defined location in the genome (14). These days, programmable nucleases are becoming a supper method to create a targeted mutation in crops which could in turn serve as a platform for molecular breeding (15). Therefore, the purpose of this review is to assess the site-directed mutation creation methods and their potential for crop improvement research.

### Site-directed mutagenesis: Basics and principles

Site-directed mutagenesis has been used to generate mutation at a single site or multiple sites of the genome (16). So far three methods of Site-directed mutagenesis are known vis. vector-based, PCR based, and Nucleases based site-directed mutagenesis. In the vector-based mutagenesis, either a plasmid or phage vector could be used for the purpose (17). In this method of mutation, one mutagenic primer and one normal primer could be used (18, 19). In the PCR based site-directed mutagenesis, the mutation could occur on double-stranded DNA and the procedure is involving simultaneous annealing of two oligonucleotide primers one mutagenic and the other normal primer annealed to the denatured double-stranded DNA (20). The nucleases based site-directed mutagenesis involves enzymes that cut DNA at a specific sequence.

Site-specific recombinase enzymes catalyze double-stranded DNA exchange between strands which have a limited degree of sequence homology. These enzymes attach to the recognition site, which is between 30 to 200 nucleotide lengths and cleave the DNA backbone (21). The *cre-lox* system consists of two components derived from the P1 bacteriophage, the *Cre* recombinase, and the *loxP* recognition site. The P1 bacteriophage uses these components as part of the natural viral life cycle and researchers have adapted the components for gene manipulation purposes (22, 23).

Transcription Activator-Like Effector Nuclease (TALEN) is another engineered nuclease, which shows a better specificity and efficiency than ZFN. Similar to ZFNs, TALENs use DNA binding motifs to direct the same non-specific nuclease to cleave the genome at a specific site, but instead of recognizing DNA triplets, each domain recognizes a single nucleotide (Table 1) (24). Clustered regularly interspaced short palindromic repeats (CRISPR technology) are the latest exciting development in gene-editing technology. The CRISPR system is RNA based bacterial defense mechanisms designed to recognize and eliminate foreign DNA from invading bacteriophages and plasmids (25).

**Table 1.** Summary of Enzymes used in Site-Directed Mutagenesis

| Feature | ZFNs | TALENs | CRISPR-Cas9 | Site-specific Recombinase |
| --- | --- | --- | --- | --- |
| Sources of enzymes | Found in Bacteria and Eukaryotes<br>zinc-binding domains in the protein transcription factor IIIA from *Xenopus* oocytes.<br>DNA Cleavage Domain FokI type II restriction enzyme Naturally found in *Flavobacterium* | Eukaryotes | Bacteria<br>(*Streptococcus* sp.) | Example Cre loxP, It is found from prokaryotes and lower eukaryotes<br>CreloxP is made by integration of bacteriophage lambda and connected with the bacterial chromosome |
| Length of recognized DNA target | 9–18 bp | 30–40 bp | 22 bp + PAM sequence | From a few hundred to tens of thousands of nucleotide pairs |
| Mechanism of target DNA recognition | DNA–protein interaction | DNA–protein interaction | DNA–RNA interaction via Watson-Crick base pairing | Site-specific recombination systems mediate DNA rearrangements by breaking and joining DNA molecules at two specific sites, termed recombination targets |
| Mechanism of DNA cleavage and repair | Double-strand break induced by *FokI* | Double-strand break induced by *FokI* | Single- or double-strand break induced by Cas9 | Double-stranded break induced by Cre gene enzyme/recombinase |
| Binding specificity | 3 Nucleotides | 1 Nucleotide | 1:1 Nucleotide pairing | To the spacer of the 13bp repeats |
| Mutation rate (%) | 10 | 20 | 20 | From hundreds to thousands of bps |
| Target site length (bp) | 18–36 | 24–40 | 22 | With an average mutation rate of ~11 amino acid substitutions per variant |
| Double-stranded break pattern | Staggered cut (4–5 nt, 5' overhang) | Staggered cut (Heterogeneous overhangs) | Sp Cas9 creates blunt ends; Cpf1 creates a staggered cut (5' | Cleaves double-stranded between 13bp repeats of loxP |
|  |  |  | overhang) |  |
| Challenges of the technology | High Off-target effects<br>Difficult to design<br>It is time consuming compared to other tools<br>Costly technology<br>FokI fusing protein unlike CRISPR Cas9 | Low<br>Moderate to design | Variable<br>Easy to design | May not be effective if the Cre gene and the loxP are separately introduced |
| Dimerization required | Yes | Yes | No | No |
| Best suited for | Gene knockout, transcriptional regulation | Gene knockout, transcriptional regulation | Gene knockout, transcriptional regulation, base editing | Gene knock out |

### Application of site-directed mutagenesis in crop improvement

Feeding the growing human population is an increasingly difficult task, and an important part of the solution acquired for the development of improved new crop cultivars with high yield and stress tolerance. As we are facing challenges to increase global agricultural productivity, there is a rapid need to accelerate the development of these traits in crops (9). Among the multitude of approaches that are used in crop improvement, targeted gene mutation technologies using different mutagenic tools are attractive technology to develop novel traits (26–28).

In recent times, different alternatives are used to bring targeted mutation and producing economic traits in crops or eliminates bad crop traits (25, 29). The rapid development of the field has allowed the development of highly efficient, precise, and cost-effective means to develop improved crop mutants (30). It has been applied and improved the yield, quality, stress tolerance traits of major food crops including maize, rice, wheat, barley, potato, soybean, carrot, cabbage, and tomato and became an appeal to molecular breeding.

### Maize improvement using site-directed mutagenesis

Maize is a major crop used as food in most of the world. Several research activities have been done to achieve targeted mutations on maize using different site-directed mutagenesis tools (31, 32).

Using targeted mutagenesis on a conserved lysine residue, Lys was replaced by Asn, Glu, or Arg to improve phosphor enolpyruvate enzyme catalytic efficiency and regulatory role on maize using plasmid vectors. As a result, the maximum velocity (Vmax) of the enzyme decreased to 22% when Asn was replaced in place of Lys and 2%, and Vmax reduction was observed when Lys is replaced by Glu in PEPC enzyme catalytic efficiency thereby enhancing sugar production (33). This indicates the potential of site-directed mutagenesis to improve affinity and catalytic efficiency of PEPC for crop physiology. In a research report when Thr was substituted by Ser using double-stranded plasmid vector site-directed mutagenesis, the regulatory capacity of Pyruvate orthophosphate Dikinase (PPDK) enzyme in maize, got improved while catalytic efficiency remains unchanged (34).

Selectable marker gene elimination from a transformed maize was reported using Cre/loxP specific recombination system with the removal of yellow fluorescent protein (yfp), which was used as a selectable marker (35). This technology could be potentially used for efficient removal of other selectable markers from genetically modified crops. In another report, site-directed mutagenesis using engineered endonuclease and targeting Lig34-site in the vicinity of the LIGULELESS1 (LG1), induction in the large-scale experiment produced 718 Parental (T_0_) plants. Altogether, the 781 T_0_ transgenic plants were evaluated by PCR, and 23 T_0_ plants were identified that contained mutations at the LIGULELESS1 locus based on visual screening of the Lig34-site (36).

Another work using I-Cel, homing endonuclease enzyme mutation was made at ms26 genomic site of maize produced small deletion and insertion of EMS26 (fertility gene at chr 5 with 5 exons) 22bp targeted site and the T_0_ maize plants carried mutated alleles of *MS26* gene which made the maize male sterile, this favors cross-breeding of the crop (37). Targeted site-directed mutation performed on five gene regions of maize (EMS26 and MS45 fertility genes, *ALS1* and *ALS2* acetolactate synthase genes, and Linguless1/*LIGI* gene) using CRISPR/Cas9 system, the mutation occurred at *ALS2* and make the crop chlorsulfuron herbicide-tolerant embryo regeneration. Moreover, targeted mutations from *EMS26* and *MS45* genes produced sterile male maize even at doubled transformation efficiency than done by engineered endonuclease (38). Stable knockout of the phytoene synthase *PSY1* gene from maize using the CRISPR/Cas9 system has been reported. The gene knockout increases sugar synthesis and whitens the powder which in turn increases market value (39). By another recent research work, sterile male maize was developed using the CRISPR/Cas9 system on zmtms5 gene (chr 2 with exon number 5) mutation and produced thermo-sensitive (32°C) maize. This is important for out-crossing and to produce improved hybrid seed (40).

An increment in maize grain yield under drought-stressed condition was reported by changing the promoter of the *ARGOS8* gene (found at chr 5 with exon number 3). *ARGOS8* gene is a negative regulator of ethylene response and modulates ethylene signal transduction and enhances drought tolerance by reducing leaf size and grain yield development. The deletion of 550-bp at genomic DNA fragment between CTS3 and CTSI removes the part of *ARGOS8 5’ UTR and* the upstream promoter sequence and native maize *GOS2* promoter was used to replace the native promoter of gene *ARGOS8* using CRISPR/Cas9 system, the protein which suppresses ethylene production was performed (41).

TALENs and CRISPR/Cas9 systems were evaluated for their targeted mutation creation efficiency at a specific site from the maize protoplast genome. Both of the tools achieved a similar mutation rate (13.1%) (Table 2). These tools could be used alternatively for maize genome editing (31). In another research using TALENs, a 10% targeted mutation rate was reported (Table 2), and it was proved that the mutation could pass to the next generations (42). Targeted mutation of Argonate18 (*zmAgo18a* and *zmAgo* 18b) and dihydroflavinol-four reductase maize genes (a1 and a4) using CRISPR/Cas9 technology resulted in a 70% mutation rate and was proved that the targeted mutation could pass to the next generations (43). From different research reports on maize site-directed mutagenesis using vector method and gene editing mutagenesis, the TALENs and CRISPR/Cas 9 system are found to be promising and repeatedly used tools to improve maize traits (31, 41, 43).

**Table 2.**
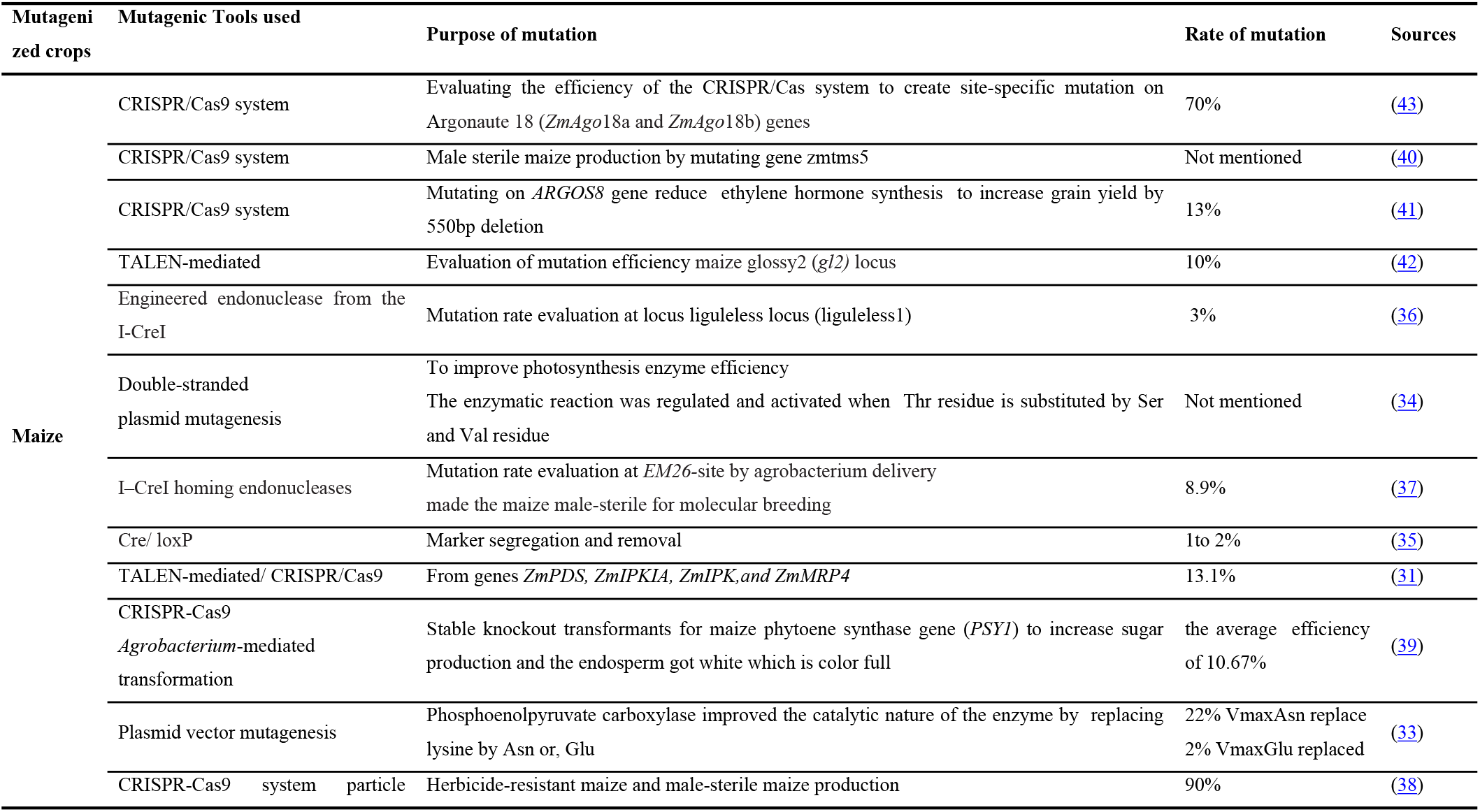
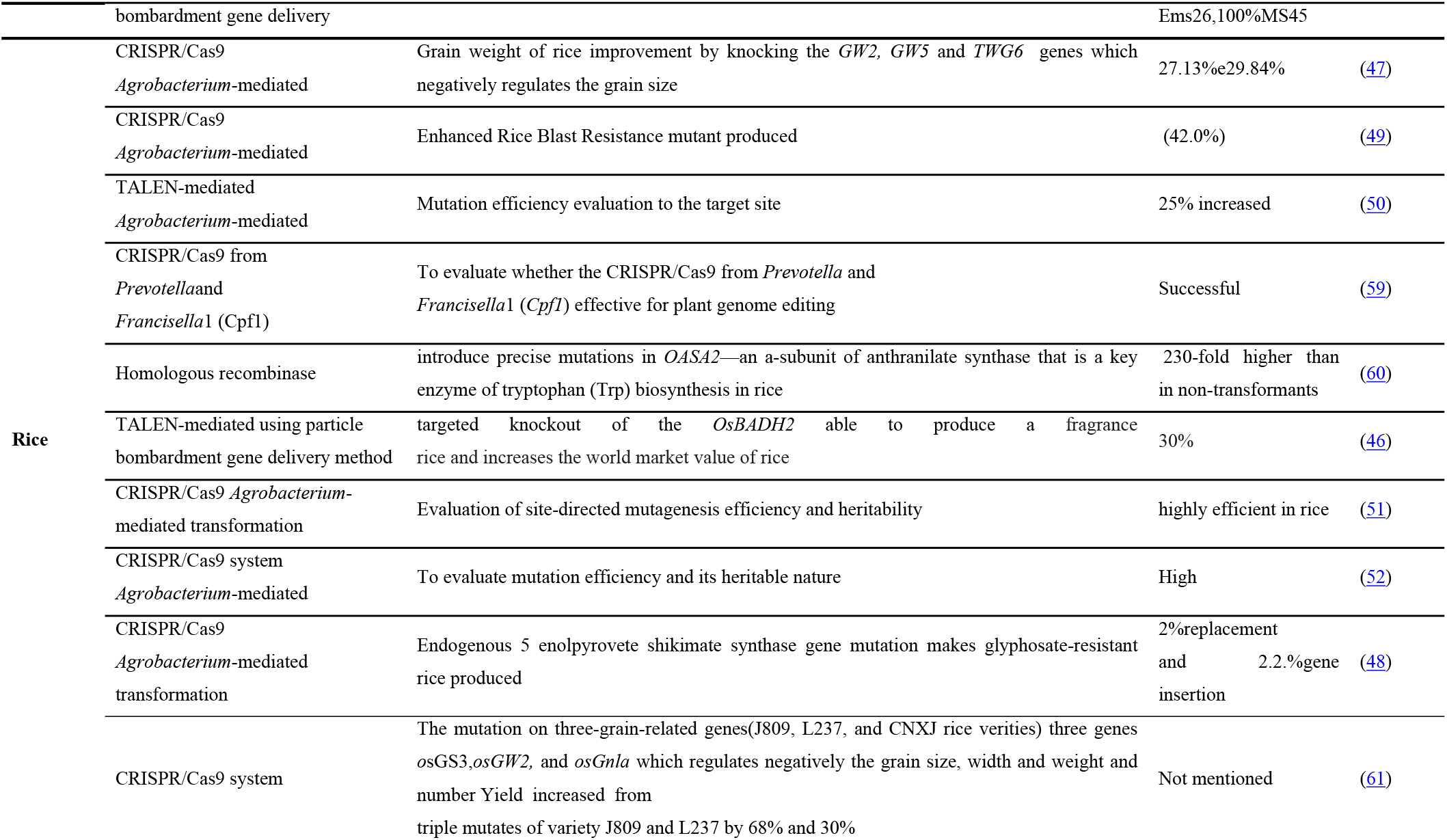
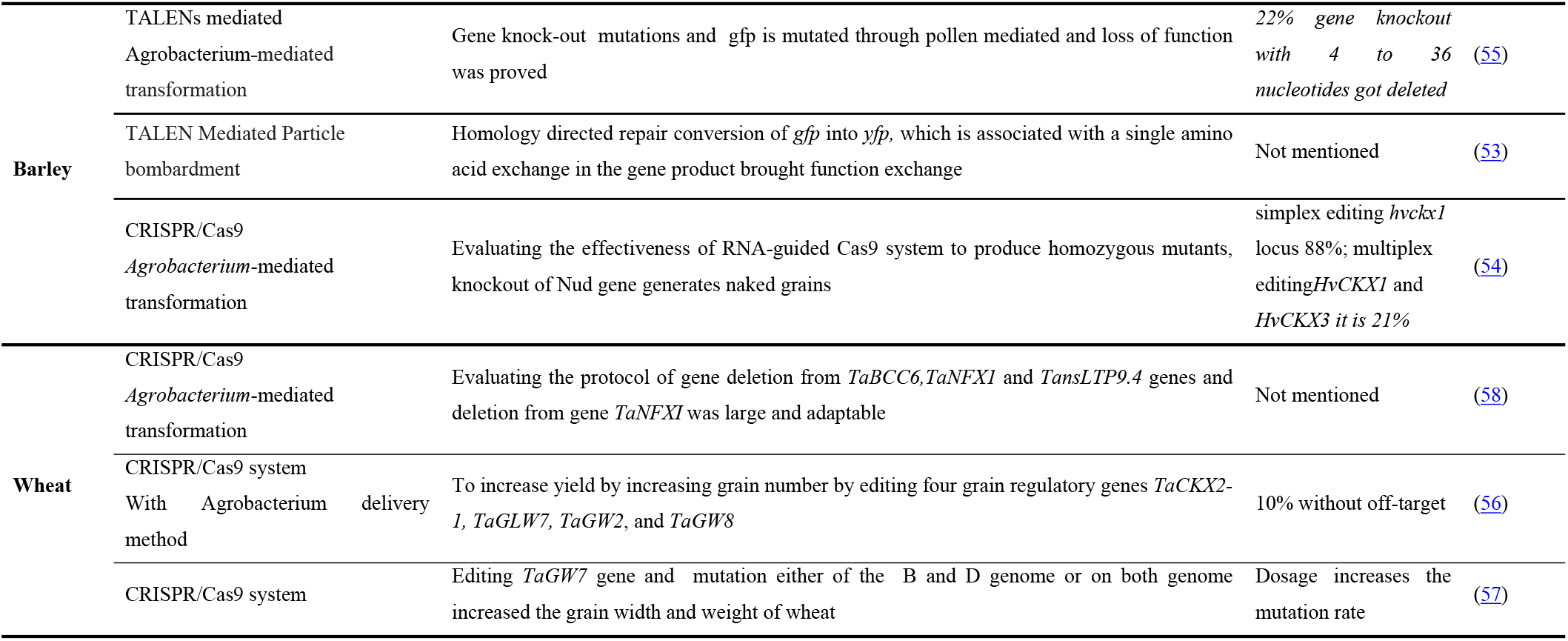
Cereals crops which have been mutagenized using different technologies

### 3.2 Rice improvement using site-directed mutagenesis

High tryptophan rice was established using a vector method of site-directed mutagenesis. The introduction of *S126F, Y367A*, and *L530D* mutations into *OASA2* was performed using a Quick Change II XL site-directed mutagenesis kit (Strata gene). Interestingly, mature seeds of homozygous GT plants accumulated Tryptophan (Trp) levels 230-folds higher than the non-transformants without any apparent morphological developmental change (Table 2) (44). Thus, from this advanced work, the great potential nutritional benefit for both humans and livestock that could not be achieved by conventional mutation was succeeded in direct crop improvement using site-directed mutagenesis.

Fragrance increases the marketability of rice (45). The development of high fragrant rice by knocking of *osBADH2* gene (found at chr 5 with 15 exon number) which produces Betain aldehyde dehydrogenase was reported using TALENs technology. TALENs were engineered to target and disrupt the *OsBADH2* gene and a total of six T_0_ heterozygous mutants *BADH2* rice plants (badh2-1 tobadh2-6) were recovered from 20 transgenic hygromycin-resistant plants. Plants badh2-2 and badh2-5 with 1-bp and 10-bp deletions, respectively, caused frameshifts at the fourth exon position and inactivated the gene and favored the biosynthesis of 2-acetyl-1-pyrroline (2AP) (46).

In recent research works, site-directed mutagenesis using the CRISPR/Cas9 system has been reported on important traits in rice like grain weight improvement, glyphosate-resistant, and blast-resistant rice development (47–49). Rice grain weight improvement by gene knockout using the CRISPR/Cas9 system was reported and among eight-grain weight controlling genes, a mutation on *GW2, GW5*, and *TGW6* genes brought weight gain for rice (47). In another report, glyphosate-resistant rice has been developed by intron mediated site-specific gene replacement and/or insertion using the CRISPR/Cas9 system. Gene replacement in the rice endogenous gene 5-enolpyruvate shikimate synthase (EPSPS) at a frequency of 2% and gene insertion at a frequency of 2.2% rice harboring the *osEPSPS* gene with intended substitutions found to be glyphosate-resistant (48). Blast resistant rice has been developed by targeted mutation using CRISPR/Cas9 SSN (C-ERF922) targeted mutation on the *osERF922* gene (49). During the targeted mutagenesis among 50 T_0_ plants 21 plants were with targeted mutation (42%) which was blast-resistant rice.

Moreover, site-directed mutagenesis efficiency and heritability also assessed by different scholars using TALENs and CRISPR/Cas9 systems (50–52). In research work using the CRISPR/Cas9 system, 11 rice genes were mutated to which the mutation rate was found to be high and heritable, but the result of mutations was not important for the agricultural development of rice (52). In another report, rice gene mutation using the CRISPR/Cas9 system ranging from 2% to 16%, which was proved to pass to the next generations (51). Development of sterile male rice enhanced grain yield, and drought-tolerant rice has been achieved by targeted mutation using TALENs. Besides the evaluation of mutation rate on targeted genes and the passage of mutant traits to subsequent generations. The target genes were *osCSA, osPMS3, osDERF1, osGN1a, osJAD1, osMST7*, and *osMST8* and the mutation of on *osCSA* and *osPMS3* resulted in photoperiod sensitive male sterility which was used after hybrid seed production and mutation on *osGN1A* and *osDERF1* generated enhanced grain yield (Table 2) and drought resistance respectively (50).

### Barely Improvement Using Site-Directed Mutagenesis

Few research works have reported, at different times by different researchers, evaluated the efficiency of site-directed mutagenesis and transmission to the next generations using TALEN and CRISPR/Cas9 system. All the research works reported that site-directed mutagenesis was efficient and was transmitted to the next T1 generations (53–55). The first transformation was made using TALEN and gene knock out through pollen regenerable cells to establish the generation of true breeding of barley. A *gfp* specific TALENs via Agrobacterium-mediated gene delivery and 22% homologous primary mutants proved to knock out of *gfp* gene, loss of function the deletion of nucleotides between 4 and 36pb length (55). By another researcher work using TALENs targeted to gfp gene with the single amino acid change, *yfp* is produced using site-specific mutation(53). Barely has been modified and the highest mutation rate was reported in simplex editing of the cytokinin oxidase/dehydrogenase *HvCKX1*) gene with a mutation rate 88% of the screened T_0_ plants (Table2) using CRISPR/Cas9 system. Multiplex editing of two genes *HvCKX1* and *HvCKX3* by obtaining 9 plants (21%) of all edited plants. The knockout of the *Nud* gene produces naked barley grains which are detectable phenotypically. Such grain reduces farming processing. It was also proven the mutation transmitted to the next generation T1 (54).

### Wheat improvement using site-directed mutagenesis

A couple of independent research works were reported on wheat grain yield-related trait improvement by targeted mutation using CRISPR/Cas9 systems for gene editing (56, 57). Targeted mutation on four grain negatively regulating genes (*TaCKX2-1, TaGLW7, TaGW2*, and *TaGW8*) and homozygous for 1160bp deletion in *TaCKX2-D1* wheat gene significantly increased grain number per spikelet (56). Edition on gene *TaGW7* to silent its expression and mutation on either B or D genome or both genomes increased both grain width and grain weight of wheat. The wheat traits that double-copy mutants showed larger yield improvement than single copy mutants (57). Using CRISPR/Cas9 system for targeted mutation produced site-specific deletion and the protocol was targeting three genes, *TaABCC6, TaNFX1*, and *TansLTP9.4* in a wheat protoplast assay and the deletion has occurred from the two genes amongst the three genes and the edit on gene *TaNFXL1* with larger deletion found to be successful (Table 2) and adaptable (58).

### Potato improvement using site-directed mutagenesis

Cold storage of potato tubers is mostly used to reduce sprouting and extending postharvest shelf life (62). However, cold temperature stimulates the reduction of sugar accumulation in potato tubers (63). In this regard, research work has been reported on potato post-harvest processing improvement using TALENs to knockout *VInv* gene within the commercial potato variety. From 600 regenerated plants, 18 plants showed mutation at least one *VInv* gene and five of these plants had mutations in all *VInv* genes. Tubers with full *VInv* gene knock out (Table 3) plants showed a noticeable level of reducing sugars, and processed chips contained reduced levels of acrylamide and were light-colored. Moreover, seven of the transformed, out of 18 modified, plant lines appeared to contain no TALEN DNA insertion in the potato genome (64). This research work could be crucial to reduce the post-harvest loss of potato and plays a role in food sustainability programs.

**Table 3.**
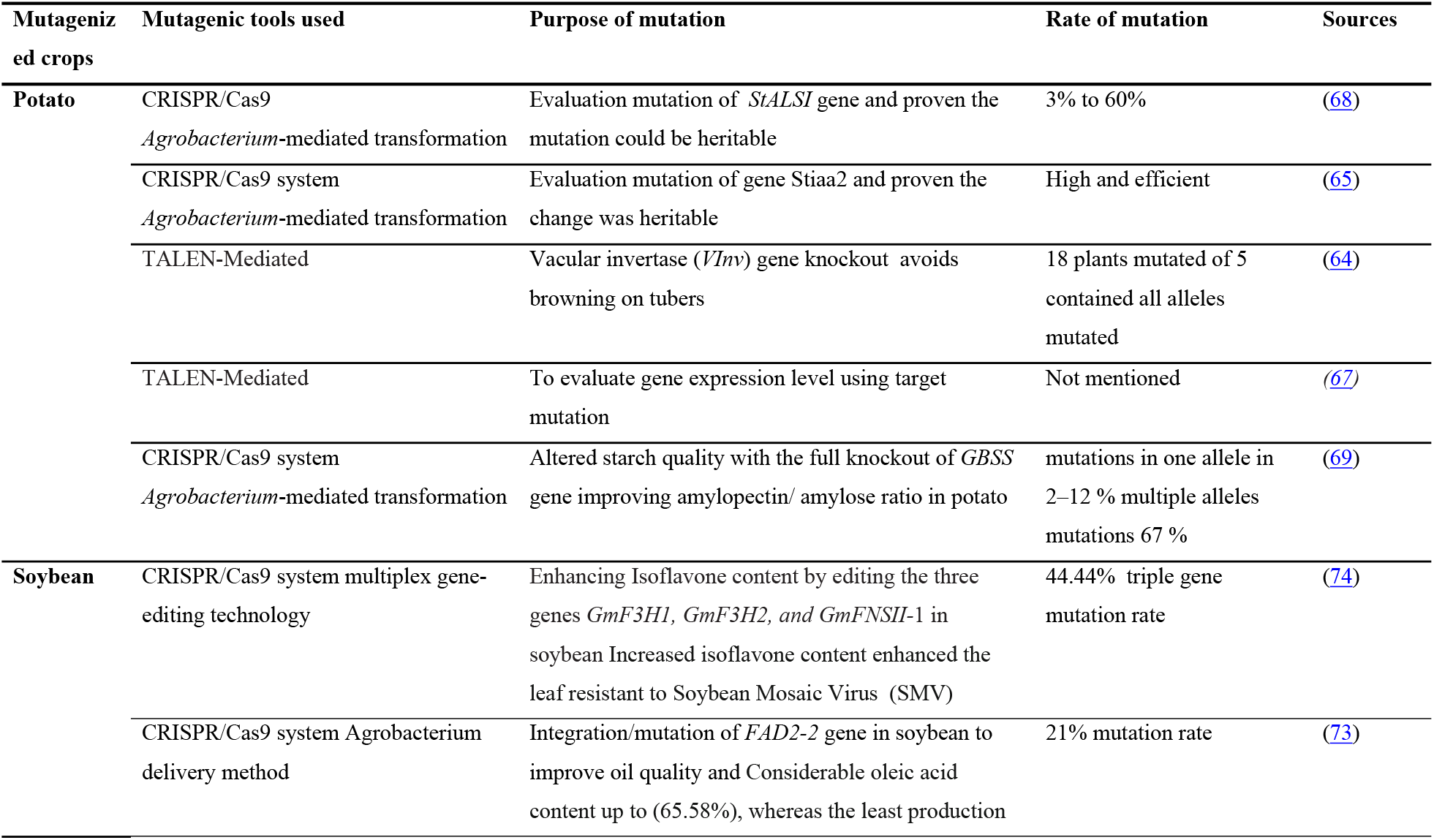
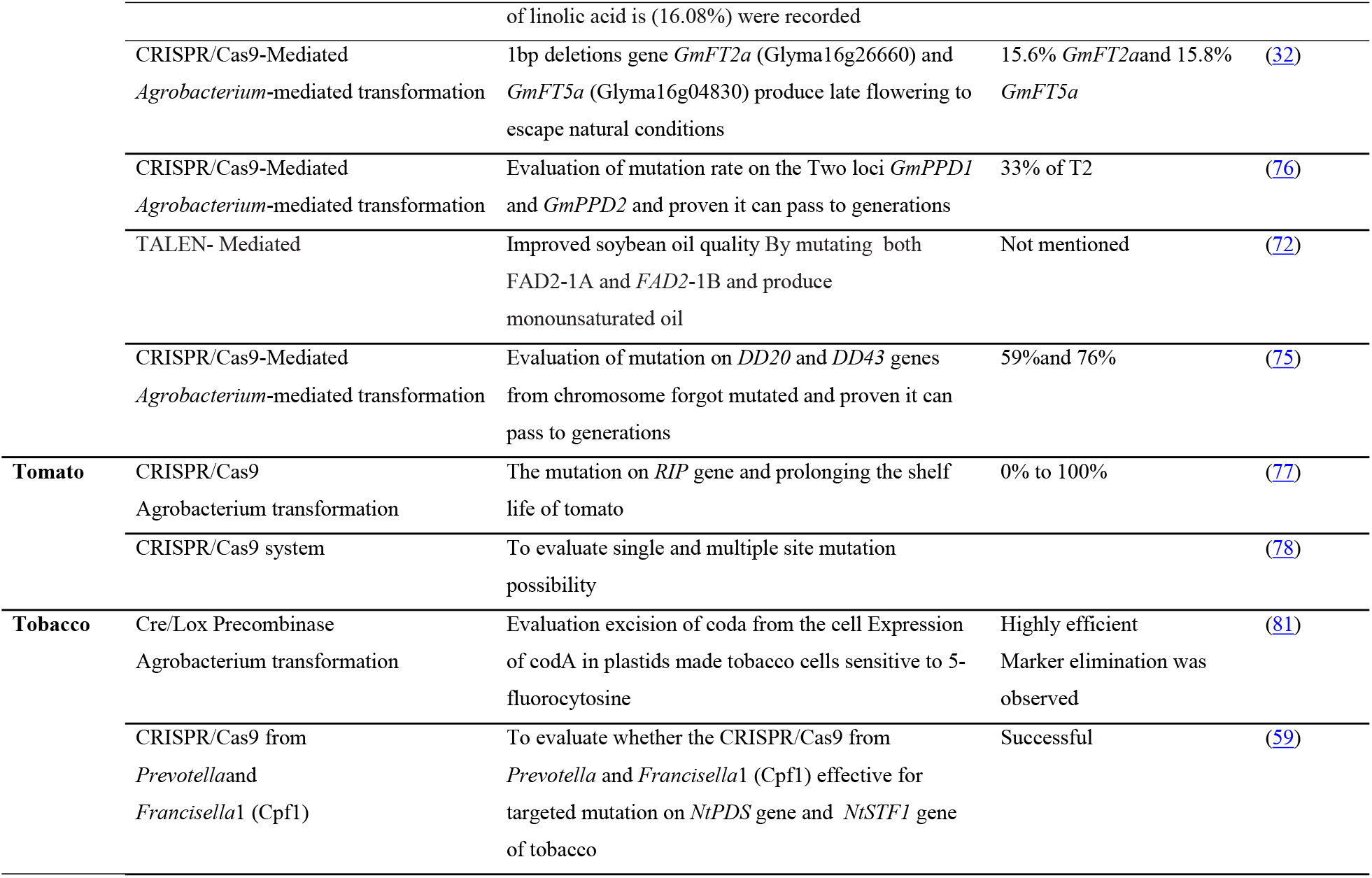
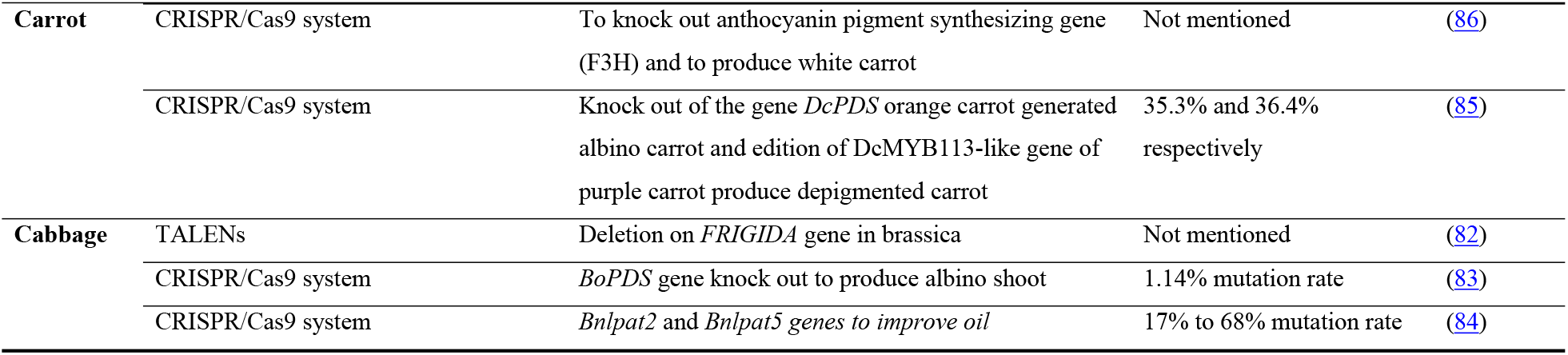
Horticultural crops, oilseed crops, and drug crops which have been mutagenized using different technologies

| **Mutagenized crops** | **Mutagenic tools used** | **Purpose of mutation** | **Rate of mutation** | **Sources** |
| --- | --- | --- | --- | --- |
| **Potato** | CRISPR/Cas9<br>*Agrobacterium*-mediated transformation | Evaluation mutation of *StALSI* gene and proven the mutation could be heritable | 3% to 60% | (68) |
|  | CRISPR/Cas9 system<br>*Agrobacterium*-mediated transformation | Evaluation mutation of gene *Stiaa2* and proven the change was heritable | High and efficient | (65) |
|  | TALEN-Mediated | Vacular invertase (*VInv*) gene knockout avoids browning on tubers | 18 plants mutated of 5 contained all alleles mutated | (64) |
|  | TALEN-Mediated | To evaluate gene expression level using target mutation | Not mentioned | (67) |
|  | CRISPR/Cas9 system<br>*Agrobacterium*-mediated transformation | Altered starch quality with the full knockout of *GBSS* gene improving amylopectin/ amylose ratio in potato | mutations in one allele in 2–12 % multiple alleles mutations 67 % | (69) |
| **Soybean** | CRISPR/Cas9 system multiplex gene-editing technology | Enhancing Isoflavone content by editing the three genes *GmF3H1*, *GmF3H2*, and *GmFNSII-1* in soybean Increased isoflavone content enhanced the leaf resistant to Soybean Mosaic Virus (SMV) | 44.44% triple gene mutation rate | (74) |
|  | CRISPR/Cas9 system *Agrobacterium* delivery method | Integration/mutation of *FAD2-2* gene in soybean to improve oil quality and Considerable oleic acid content up to (65.58%), whereas the least production of linolic acid is (16.08%) were recorded | 21% mutation rate | (73) |
|  | CRISPR/Cas9-Mediated<br>*Agrobacterium*-mediated transformation | 1bp deletions gene *GmFT2a* (Glyma16g26660) and *GmFT5a* (Glyma16g04830) produce late flowering to escape natural conditions | 15.6% *GmFT2a* and 15.8% *GmFT5a* | (32) |
|  | CRISPR/Cas9-Mediated<br>*Agrobacterium*-mediated transformation | Evaluation of mutation rate on the Two loci *GmPPD1* and *GmPPD2* and proven it can pass to generations | 33% of T2 | (76) |
|  | TALEN- Mediated | Improved soybean oil quality By mutating both *FAD2-1A* and *FAD2-1B* and produce monounsaturated oil | Not mentioned | (72) |
|  | CRISPR/Cas9-Mediated<br>*Agrobacterium*-mediated transformation | Evaluation of mutation on *DD20* and *DD43* genes from chromosome forgot mutated and proven it can pass to generations | 59% and 76% | (75) |
| Tomato | CRISPR/Cas9<br>*Agrobacterium* transformation | The mutation on *RIP* gene and prolonging the shelf life of tomato | 0% to 100% | (77) |
|  | CRISPR/Cas9 system | To evaluate single and multiple site mutation possibility |  | (78) |
| Tobacco | Cre/Lox Precombinase<br>*Agrobacterium* transformation | Evaluation excision of coda from the cell Expression of *codA* in plastids made tobacco cells sensitive to 5-fluorocytosine | Highly efficient<br>Marker elimination was observed | (81) |
|  | CRISPR/Cas9 from<br>*Prevotella* and<br>*Francisella*1 (Cpf1) | To evaluate whether the CRISPR/Cas9 from *Prevotella* and *Francisella*1 (Cpf1) effective for targeted mutation on *NtPDS* gene and *NtSTF1* gene of tobacco | Successful | (59) |
| **Carrot** | CRISPR/Cas9 system | To knock out anthocyanin pigment synthesizing gene (F3H) and to produce white carrot | Not mentioned | (86) |
|  | CRISPR/Cas9 system | Knock out of the gene *DcPDS* orange carrot generated albino carrot and edition of *DcMYB113*-like gene of purple carrot produce depigmented carrot | 35.3% and 36.4% respectively | (85) |
| **Cabbage** | TALENs | Deletion on *FRIGIDA* gene in brassica | Not mentioned | (82) |
|  | CRISPR/Cas9 system | *BoPDS* gene knock out to produce albino shoot | 1.14% mutation rate | (83) |
|  | CRISPR/Cas9 system | *Bnlpat2* and *Bnlpat5* genes to improve oil | 17% to 68% mutation rate | (84) |

Research works were reported on the evaluation of mutation efficiency and the heritability of the mutation created by CRISPR/Cas9 system and TALEN to the subsequent generations (65–67). The research work using the CRISPR/Cas9 system for targeted mutation on the *stALS1* gene reported mutation ranging from 3% to 60%, and the mutation was proved its heritability to the next generation (68). A site-directed mutation on the *stIAA2* gene using CRISPR/Cas9 system resulted a high and efficient mutation, and the change was proved as heritable to the next potato generation (65).

Using TALENs, site-directed mutagenesis on the *stALS* gene resulted in a higher mutation rate (Table 3) that was proven to be transferred to the next generations (67). Starch quality was altered using site-directed mutagenesis on the *GBSS* gene function using CRISPR/Cas9 technology. In this work, the *GBSS* gene has been fully knocked out in the protoplast of tetraploid potato, and mutation was produced in all four alleles. At three regions of the gene granule bound starch synthase were targeted and resulted in mutation at least one allele in 2% to 12% regenerated shoots and multiplex mutation was up to 67%. The removal of *GBSS* enzyme activity leads to starch with altered amylose synthesis concomitant increase in the amylopectin/amylose ratio (69).

### Soybean improvement using site-directed mutagenesis

Soybean oil quality improvement has been reported by targeted mutagenesis of the fatty acid desaturase two gene families (*FAD2-1A and FAD2-1B*) using TALENs. The desaturase removes hydrogen from fatty acids and makes the poly unsaturation which could be a threat to heart and brain health (70). The trans-fatty acids produced through hydrogenation pose a health threat (71). Four of the 19 transgenic soybean line mutations in both *FAD2-1A* and *FAD2-1B* were observed in DNA taken from leaf samples. The fatty acid from homozygous mutant seeds of *FAD2-1A* and *FAD2-1B* oleic acid which is a monounsaturated fatty acid (18:1 cis-9) which was omega fatty acid increased from 20% to 80% and linoleic acid polyunsaturated fatty acid (omega 6 fatty acids) decreased from 50% to 4% (Table 3) and the mutation has proven as heritable (72). Another research on soybean oil improvement using CRISPR/Cas 9 system for editing *FAD2-2* soybean gene reported a 21% mutation rate with improved oil quality. A considerable oleic acid content (up to 65.58%), and the least production of linolic acid (16.08%) was recorded (73). Hence, site-directed mutagenesis through gene editing could be one potential for nutritional improvement in food crops.

Recent research work on adaptable soybean to climate change by altering the flowering time of soybean improved by targeted mutagenesis using the CRISPR/Cas9 system. Cultivar Jack was mutated at the specific site and T1-generation soya bean plants homozygous for null alleles of GmFT2a (chr.16 with four exon number) frameshift mutated by a 1-bp insertion or short deletion resulted knocking of the gene thereby producing a trait, late-flowering period to escape the natural condition to adapt the stress and the mutation was proved as it is heritable (32). Soybean nutritional improvement and viral disease tolerant have been reported using gene editing targeted mutagenesis. Multiplex gene editing using the CRISPR/Cas9 system on three genes (*GmF3H1, GmF3H2, and GmFNSII-1*) in soybean which had negative regulation of isoflavone production has been knocked out. The triple gene mutation efficiency was 44.44%. The T_3_ homologous triple gene mutants increased Isoflavone content in the leaf twice and the crop becomes resistant to soybean mosaic virus due to the increased isoflavone metabolite (74).

The mutation rate and heritability of directed mutagenesis using the CRISPR/Cas9 system with Agrobacterium-mediated transformation of soybean were reported (75, 76). Two genomic sites of soybean DD20 and DD43 mutagenized using the CRISPR/Cas9 system and it was reported with a mutation frequency of 59% and 76% respectively and the mutation was proven as heritable to the next T1 generations (75). Simultaneous site-directed mutagenesis of *GmPPD* loci using CRISPR/Cas9-system soybean mutation in *GmPPD* confirmed 33% of the T2 seeds and it was proven that the mutation was heritable (76). Among six research works reviewed on soybean site-directed mutagenesis, five of the works were done using CRISPR/Cas9 technology which was efficient to generate targeted mutation.

### Tomato improvement using site-directed mutagenesis

Tomato shelf life has been improved by site-directed mutagenesis using the CRISPR/Cas9 system by the agrobacterium gene delivery method. Gene deletion/insertion on the *RIN* gene which encodes a MAD1-box transcription factor regulating fruit ripening. The *RIN* protein defective mutants were found to be effective to make the tomato stay fresh for several months by changing the ripening physiology and ethylene production (77). The mutation rate also ranges from 0% to 100%. Targeted mutation employed by CRISPR/Cas9 system as using the Agrobacterium-mediated gene delivery method and single and multi-site mutagenesis has been reported and summarized as this technology could be employed to produce site-directed mutagenesis on important traits from the same or other crops (78).

### Tobacco crop site-directed mutagenesis

Incorporation of the selectable marker gene during plant transformation is crucial to know the whereabouts of the gene of interest (79). However, the selectable marker gene especially the old markers remain a public concern (80). Using *CRE-lox* recombinase *CoDA* selectable marker flanked with two directly oriented lox sites, the highly efficient elimination of the marker gene (Table 3) introduced through pollination was reported (81). By another research report, depigmentation and leaf size expansion has been done by a targeted mutation on *NtPDS* and *NsSTF1* genes respectively by using the CRISPR/Cas 9 system. The CRISPR/Cas9 was optimized with *FnCpf1* with a 24 nt targeted sequence that was optimized to induce mutation in two genes. These genes are phytoene desaturase (*NtPDS*) and STENOFOLLA ortholog in *Nicotiana tobacum* (NtSTF1).Mutation in the first gene resulted depigmentation of chlorophyll and albino tobacco was produced moreover mutation on the second (*NtSTF1*) gene induced tobacco leaf blade expansion and enhances production for tobacco(59).

### Cabbage (*Brasica oreracea*) improvement through site-directed mutagenesis

Cabbage is a vegetable and oilseed crop. A research work on Cabbage (*Brasica oreracea L. var*) using customized TALEN based nuclease constructed using a method “unit assembly” specially targeted the endogenous FRIGIDA gene in *Brasica oreracea L. var* modified the targeted site with deletion and this protocol is proven to bring genetic modification by site-directed mutagenesis (82). Two other research works that have been reported on a targeted mutation of the cabbage genome using the CRISPR/Cas9 system (83, 84). One report has been done on *BoPDS* gene knockout and produced albino cabbage (*Brassica oleracea*) shoot and the mutation rate was 1.14% (83). By another work report Knockout of *Bnlpat2* and *Bnlpat5* genes have been done with mutation rate ranging 17 to 68% to improve the seed starch content and increased oil bodies of the matured seed of cabbage (84). From these two reports no off-target edition hence the CRISPR/Cas 9 system mutagenesis is potential and efficient for crop improvement.

### Carrot improvement through site-directed mutagenesis

Two independent research works reported improvement crops by site-directed mutagenesis using the CRISPR/Cas 9 system in 2019 (85, 86). In the first report anthocyanin pigment synthesizing gene (F3H) knock out and produced white carrot. The research output depigmented the purple color into a white color carrot and produced marketable products by mutating the F3H gene through deletion and removed anthocyanin expression (86). By the second research report knockout CRISPR/ Cas 9 technology of the gene, DcPDS orange carrot generated albino carrot with a mutation rate of 35.3%, and edition of DcMYB113-like gene of purple carrot produces depigmented carrot with a mutation rate of 36.4% (85).

These days new insight for crop improvement has been added and used as a means to improve crop of interest. New tools are discussed in different research findings and their potency to produce site-specific genetic alteration and the extent of its heritability to the proceeding generation. From this review paper, 38 original research works on crop targeted mutation has been assessed four mutagenic tools have been investigated at a different frequency, at which CRISPR/Cas9 system found to be repeatedly used to transform different crops and reported as it is an efficient technology to produce the intended specific mutation (Figure 1) (43, 87). TALENs found the second tool which was used to transform crops to bring efficiently targeted mutation and it was affirmed by most scholars as it is potent technology for crop transformation (Table 2 and 3).

**Figure 1.**
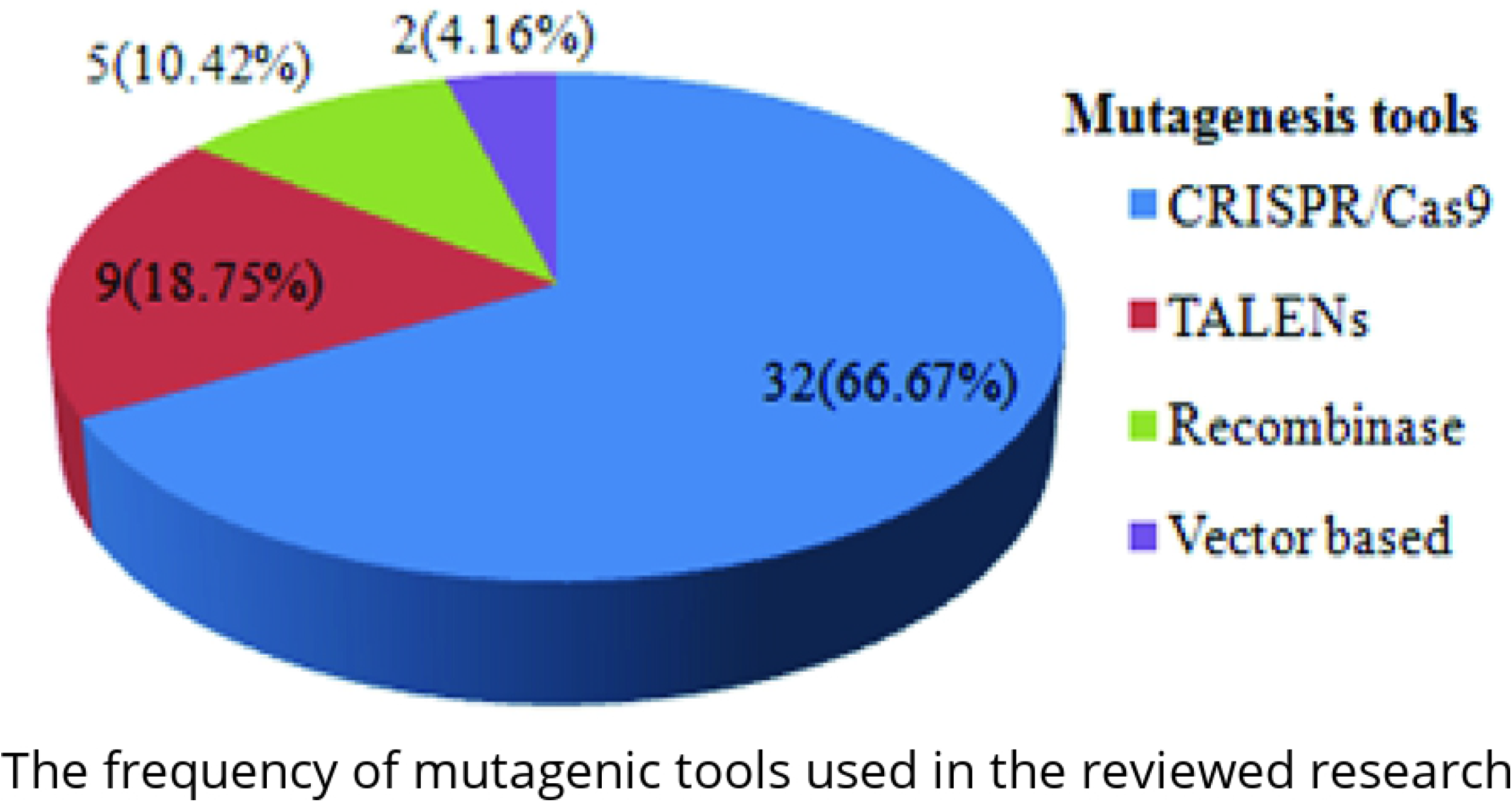
The frequency of mutagenic tools used in the reviewed research works

Furthermore, according to the research reports, these technologies were highly employed from cereal crops (Figure 2). As we know monocots are the major staple food crops to the human population around the globe (87), hence doing improvement targeting to cereals in particular and all crops at large could ensure food sustainability.

**Figure 2.**
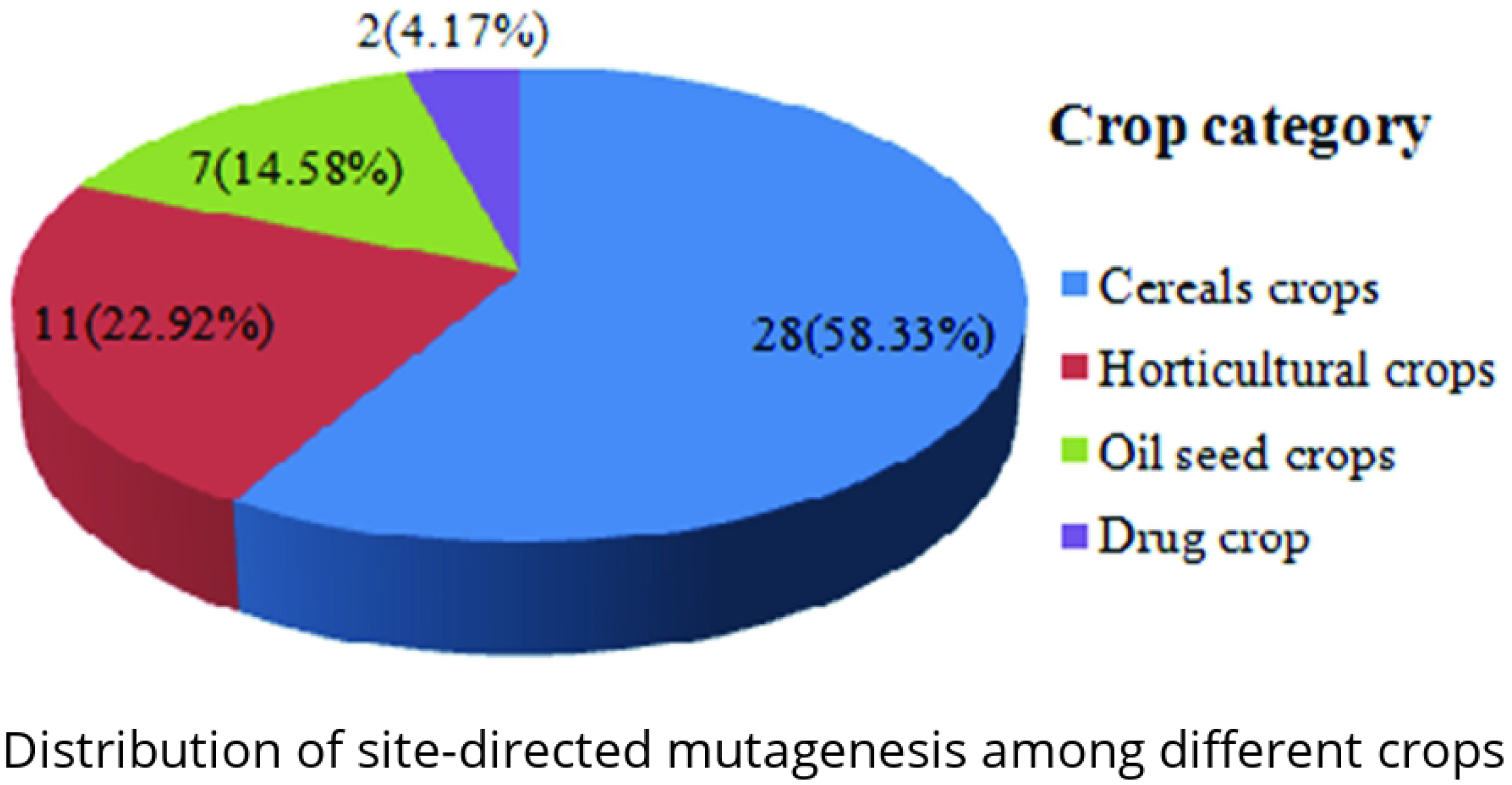
Distribution of site-directed mutagenesis among different crops groups on reviewed research works

### Challenges of site-directed mutagenesis

Site-directed mutagenesis is crucial in crop production as it is important to bring the required change on the target DNA sequence thereby changing the gene output and the trait of the crop (10). Site-specific mutagenesis is more powerful than genetic transformation through recombinant DNA technology of crops since the later introduces foreign genes to plants at a random place while the former alter the gene at the programmed site. The random integration of the introduction of the foreign gene may silent other important genes or may bring uncommon gene expressions (88, 89). All technologies developed so far might not be equally efficient in bringing the ultimate change in crop improvement using site-directed mutagenesis (87).

Developed technologies for targeted mutation creation showed advancement with time. The vector and the PCR methods of site-directed mutagenesis recently are less likely to be used for plant transformation because of their low efficiency and inadequacy in crop agronomic traits improvement (90). Hence, other new promising and highly applicable technologies have been developed and used for crop transformation through site-directed mutation (25, 28, 91, 92). Recently developed mutagenic tools also have limitations to bring efficient programmed genetic changes on a crop genome (Table 1). Zinc Finger Nucleases (ZFNs) used for site-directed mutagenesis could produce the off-target effect, large size effect to delivery, dimerization, comparatively high cost, and laborious nature.

TALENs is one of highly effective, easy to construct, and less costly tool compared to Zinc Finger Nucleases (ZFNs) to create site-specific deletion or insertion (46, 50). However, still, it has a dimerization and off-target effect to a lesser extent compared to the CRISPR/Cas9 system (82, 92). The CRISPR/Cas9 system possesses several potential advantages over ZFNs and TALENs. The short size of the sgRNA sequence makes it easier to deliver, cheap to construct, not laborious, efficient compared to others. However, this novel nucleases has still little limitations such as it experiences off-target effect (87).

### Conclusion

Crop improvement using site-directed mutagenesis employing plasmid vector based and sitespecific nucleases transformation has been summarized. From the reviewed works, CRISPR/Cas9 system was found repeatedly (66.67%) used to improve crop traits by targeted mutagenesis. TALENs were used for knockout of bad trait coding genes. Hence, the CRISPR/Case9 technology is widely used to improve the crop of their interest and to ensure food sustainability for its efficiency and less off-target effects. A lot of agronomic traits, physiological traits, and stress-tolerant traits of crops have been improved by site-directed mutation.

The site-directed mutagenesis technology is found to be highly applied for cereal crops that were less effective to be transformed using recombinant DNA technology. Cereals were the dominant crops to be transformed by site-directed mutagenesis of which maize and rice are on the front. Finally, the trend of technology usage shows that CRISPR/Cas9 technology has been highly used by researchers around the globe to bring efficient transformation and crop improvement. In addition, for knocking out of genes with bad traits, TALENs were found to be ideal. Hence, targeting and doing improvement on the major stable food crops could ensure the world’s food security.

## References

1. Mohan Jain S, Suprasanna P. Induced mutations for enhancing nutrition and food production. Gene Conserve. 2011;10(41).

2. Nestel P, Bouis HE, Meenakshi J, Pfeiffer W. Biofortification of staple food crops. The Journal of nutrition. 2006;136(4):1064–7.

3. Godfray HCJ, Beddington JR, Crute IR, Haddad L, Lawrence D, Muir JF, et al. Food security: the challenge of feeding 9 billion people. science. 2010;327(5967):812–8.

4. Rashid B, Tariq M, Khalid A, Shams F, Ali Q, Ashraf F, et al. Crop improvement: New approaches and modern techniques. Plant Gene and Trait. 2017;8(3).

5. Rao M, Rao VR, Angadi S. Crop Improvement in the World-Past, Present and Future. Asian Agri-History. 2018;22(4).

6. Caligari PD, Forster BP. Plant breeding and crop improvement. eLS. 2001:1–11.

7. Dobermann A, Nelson R. Opportunities and solutions for sustainable food production. Sustainable Development Solutions Network: Paris, France. 2013.

8. Roychowdhury R, Tah J. Mutagenesis—A potential approach for crop improvement. Crop Improvement: Springer; 2013. p. 149–87.

9. Sauer NJ, Mozoruk J, Miller RB, Warburg ZJ, Walker KA, Beetham PR, et al. Oligonucleotide-directed mutagenesis for precision gene editing. Plant biotechnology journal. 2016;14(2):496–502.

10. Oladosu Y, Rafii MY, Abdullah N, Hussin G, Ramli A, Rahim HA, et al. Principle and application of plant mutagenesis in crop improvement: a review. Biotechnology & Biotechnological Equipment. 2016;30(1):1–16.

11. Kharkwal M, Shu Q. The role of induced mutations in world food security. Induced plant mutations in the genomics era Food and Agriculture Organization of the United Nations, Rome. 2009:33–8.

12. Liu H, Naismith JH. An efficient one-step site-directed deletion, insertion, single and multiple-site plasmid mutagenesis protocol. BMC biotechnology. 2008;8(1):91.

13. Zheng L, Baumann U, Reymond J-L. An efficient one-step site-directed and site-saturation mutagenesis protocol. Nucleic acids research. 2004;32(14):e115–e.

14. Van de Wiel C, Schaart J, Lotz L, Smulders M. New traits in crops produced by genome editing techniques based on deletions. Plant biotechnology reports. 2017;11(1):1–8.

15. Modrzejewski D, Hartung F, Sprink T, Krause D, Kohl C, Wilhelm R. What is the available evidence for the range of applications of genome-editing as a new tool for plant trait modification and the potential occurrence of associated off-target effects: a systematic map. Environmental Evidence. 2019;8(1):27.

16. Forloni M, Liu AY, Wajapeyee N. Methods for In Vitro Mutagenesis. Cold Spring Harbor Protocols. 2019;2019(12):pdb. top097733.

17. Saboulard D, Dugas V, Jaber M, Broutin J, Souteyrand E, Sylvestre J, et al. High-throughput site-directed mutagenesis using oligonucleotides synthesized on DNA chips. Biotechniques. 2006;39(3):363–8.

18. Zoller MJ, Smith M. Oligonucleotide-directed mutagenesis: a simple method using two oligonucleotide primers and a single-stranded DNA template. Dna. 1984;3(6):479–88.

19. Smith M. Site-directed mutagenesis. Trends in Biochemical Sciences. 1982;7(12):440–2.

20. Braman J, Papworth C, Greener A. Site-directed mutagenesis using double-stranded plasmid DNA templates. In vitro mutagenesis protocols: Springer; 1996. p. 31–44.

21. Coates CJ, Kaminski JM, Summers JB, Segal DJ, Miller AD, Kolb AF. Site-directed genome modification: derivatives of DNA-modifying enzymes as targeting tools. Trends in biotechnology. 2005;23(8):407–19.

22. Lambert JM, Bongers RS, Kleerebezem M. Cre-lox-based system for multiple gene deletions and selectable-marker removal in Lactobacillus plantarum. Appl Environ Microbiol. 2007;73(4):1126–35.

23. Araki K, Araki M, Yamamura Ki. Site-directed integration of the cre gene mediated by Cre recombinase using a combination of mutant lox sites. Nucleic Acids Research. 2002;30(19):e103–e.

24. Li T, Liu B, Chen CY, Yang B. TALEN-mediated homologous recombination produces site-directed DNA base change and herbicide-resistant rice. Journal of Genetics and Genomics. 2016;43(5):297–305.

25. Gupta M, Gerard M, Padmaja SS, Sastry RK. Trends of CRISPR technology development and deployment into Agricultural Production-Consumption Systems. World Patent Information. 2020;60:101944.

26. Gaj T, Gersbach CA, Barbas III CF. ZFN, TALEN, and CRISPR/Cas-based methods for genome engineering. Trends in biotechnology. 2013;31(7):397–405.

27. Christian M, Cermak T, Doyle EL, Schmidt C, Zhang F, Hummel A, et al. Targeting DNA double-strand breaks with TAL effector nucleases. Genetics. 2010;186(2):757–61.

28. Eş I, Gavahian M, Marti-Quijal FJ, Lorenzo JM, Khaneghah AM, Tsatsanis C, et al. The application of the CRISPR-Cas9 genome editing machinery in food and agricultural science: Current status, future perspectives, and associated challenges. Biotechnology advances. 2019.

29. Ren C, Liu X, Zhang Z, Wang Y, Duan W, Li S, et al. CRISPR/Cas9-mediated efficient targeted mutagenesis in Chardonnay (Vitis vinifera L.). Scientific reports. 2016;6:32289.

30. Nishizawa-Yokoi A, Cermak T, Hoshino T, Sugimoto K, Saika H, Mori A, et al. A defect in DNA Ligase4 enhances the frequency of TALEN-mediated targeted mutagenesis in rice. Plant physiology. 2016;170(2):653–66.

31. Liang Z, Zhang K, Chen K, Gao C. Targeted mutagenesis in Zea mays using TALENs and the CRISPR/Cas system. Journal of Genetics and Genomics. 2014;41(2):63–8.

32. Cai Y, Chen L, Liu X, Guo C, Sun S, Wu C, et al. CRISPR/Cas9-mediated targeted mutagenesis of GmFT2a delays flowering time in soya bean. Plant biotechnology journal. 2018;16(1):176–85.

33. Dong L-Y, Ueno Y, Hata S, Izui K. Effects of site-directed mutagenesis of conserved Lys606 residue on catalytic and regulatory functions of maize C4-form phosphoenolpyruvate carboxylase. Plant and cell physiology. 1997;38(12):1340–5.

34. Chastain CJ, Lee ME, Moorman MA, Shameekumar P, Chollet R. Site-directed mutagenesis of maize recombinant C4-pyruvate, orthophosphate dikinase at the phosphorylatable target threonine residue. FEBS letters. 1997;413(1):169–73.

35. Djukanovic V, Orczyk W, Gao H, Sun X, Garrett N, Zhen S, et al. Gene conversion in transgenic maize plants expressing FLP/FRT and Cre/loxP site-specific recombination systems. Plant biotechnology journal. 2006;4(3):345–57.

36. Gao H, Smith J, Yang M, Jones S, Djukanovic V, Nicholson MG, et al. Heritable targeted mutagenesis in maize using a designed endonuclease. The Plant Journal. 2010;61(1):176–87.

37. Djukanovic V, Smith J, Lowe K, Yang M, Gao H, Jones S, et al. Male-sterile maize plants produced by targeted mutagenesis of the cytochrome P 450-like gene (MS 26) using a re-designed I-C reI homing endonuclease. The plant journal. 2013;76(5):888–99.

38. Svitashev S, Young JK, Schwartz C, Gao H, Falco SC, Cigan AM. Targeted mutagenesis, precise gene editing, and site-specific gene insertion in maize using Cas9 and guide RNA. Plant physiology. 2015;169(2):931–45.

39. Zhu J, Song N, Sun S, Yang W, Zhao H, Song W, et al. Efficiency and inheritance of targeted mutagenesis in maize using CRISPR-Cas9. Journal of Genetics and Genomics. 2016;43(1):25–36.

40. Li J, Zhang H, Si X, Tian Y, Chen K, Liu J, et al. Generation of thermosensitive male-sterile maize by targeted knockout of the ZmTMS5 gene. Journal of genetics and genomics= Yi chuan xue bao. 2017;44(9):465.

41. Shi J, Gao H, Wang H, Lafitte HR, Archibald RL, Yang M, et al. ARGOS 8 variants generated by CRISPR-Cas9 improve maize grain yield under field drought stress conditions. Plant biotechnology journal. 2017;15(2):207–16.

42. Char SN, Unger-Wallace E, Frame B, Briggs SA, Main M, Spalding MH, et al. Heritable site-specific mutagenesis using TALEN s in maize. Plant biotechnology journal. 2015;13(7):1002–10.

43. Char SN, Neelakandan AK, Nahampun H, Frame B, Main M, Spalding MH, et al. An Agrobacterium-delivered CRISPR/Cas9 system for high-frequency targeted mutagenesis in maize. Plant biotechnology journal. 2017;15(2):257–68.

44. Saika H, Oikawa A, Matsuda F, Onodera H, Saito K, Toki S. Application of Gene Targeting to Designed Mutation Breeding of High-Tryptophan Rice1 [W][OA]. American Society of Plant Biologists. 211;156:1269–77.

45. Ashokkumar S, Jaganathan D, Ramanathan V, Rahman H, Palaniswamy R, Kambale R, et al. Creation of novel alleles of fragrance gene OsBADH2 in rice through CRISPR/Cas9 mediated gene editing. PloS one. 2020;15(8):e0237018.

46. Shan Q, Zhang Y, Chen K, Zhang K, Gao C. Creation of fragrant rice by targeted knockout of the Os BADH 2 gene using TALEN technology. Plant biotechnology journal. 2015;13(6):791–800.

47. Xu R, Yang Y, Qin R, Li H, Qiu C, Li L, et al. Rapid improvement of grain weight via highly efficient CRISPR/Cas9-mediated multiplex genome editing in rice. Journal of genetics and genomics= Yi chuan xue bao. 2016;43(8):529.

48. Li J, Meng X, Zong Y, Chen K, Zhang H, Liu J, et al. Gene replacements and insertions in rice by intron targeting using CRISPR-Cas9. Nature plants. 2016;2(10):16139.

49. Wang F, Wang C, Liu P, Lei C, Hao W, Gao Y, et al. Enhanced rice blast resistance by CRISPR/Cas9-targeted mutagenesis of the ERF transcription factor gene OsERF922. PloS one. 2016;11(4):e0154027.

50. Zhang H, Gou F, Zhang J, Liu W, Li Q, Mao Y, et al. TALEN-mediated targeted mutagenesis produces a large variety of heritable mutations in rice. Plant biotechnology journal. 2016;14(1):186–94.

51. Zhang H, Zhang J, Wei P, Zhang B, Gou F, Feng Z, et al. The CRISPR/C as9 system produces specific and homozygous targeted gene editing in rice in one generation. Plant biotechnology journal. 2014;12(6):797–807.

52. Xu R, Li H, Qin R, Wang L, Li L, Wei P, et al. Gene targeting using the Agrobacterium tumefaciens-mediated CRISPR-Cas system in rice. Rice. 2014;7(1):5.

53. Budhagatapalli N, Rutten T, Gurushidze M, Kumlehn J, Hensel G. Targeted modification of gene function exploiting homology-directed repair of TALEN-mediated double-strand breaks in barley. G3: Genes, Genomes, Genetics. 2015;5(9):1857–63.

54. Gasparis S, Kała M, Przyborowski M, Łyżnik LA, Orczyk W, Nadolska-Orczyk A. A simple and efficient CRISPR/Cas9 platform for induction of single and multiple, heritable mutations in barley (Hordeum vulgare L.). Plant methods. 2018;14(1):111.

55. Gurushidze M, Hensel G, Hiekel S, Schedel S, Valkov V, Kumlehn J. True-breeding targeted gene knock-out in barley using designer TALE-nuclease in haploid cells. PloS one. 2014;9(3):e92046.

56. Zhang Z, Hua L, Gupta A, Tricoli D, Edwards KJ, Yang B, et al. Development of an Agrobacterium-delivered CRISPR/Cas9 system for wheat genome editing. Plant biotechnology journal. 2019;17(8):1623–35.

57. Wang W, Pan Q, Tian B, He F, Chen Y, Bai G, et al. Gene editing of the wheat homologs of TONNEAU 1-recruiting motif encoding gene affects grain shape and weight in wheat. The Plant Journal. 2019;100(2):251–64.

58. Cui X, Balcerzak M, Schernthaner J, Babic V, Datla R, Brauer EK, et al. An optimised CRISPR/Cas9 protocol to create targeted mutations in homoeologous genes and an efficient genotyping protocol to identify edited events in wheat. Plant methods. 2019;15(1):1–12.

59. Endo A, Masafumi M, Kaya H, Toki S. Efficient targeted mutagenesis of rice and tobacco genomes using Cpf1 from Francisella novicida. Scientific reports. 2016;6:38169.

60. Saika H, Oikawa A, Matsuda F, Onodera H, Saito K, Toki S. Application of gene targeting to designed mutation breeding of high-tryptophan rice. Plant physiology. 2011;156(3):1269–77.

61. Zhou J, Xin X, He Y, Chen H, Li Q, Tang X, et al. Multiplex QTL editing of grain-related genes improves yield in elite rice varieties. Plant cell reports. 2019;38(4):475–85.

62. Alamar MC, Tosetti R, Landahl S, Bermejo A, Terry LA. Assuring potato tuber quality during storage: a future perspective. Frontiers in plant science. 2017;8:2034.

63. Krause KP, Hill L, Reimholz R, Hamborg Nielsen T, Sonnewald U, Stitt M. Sucrose metabolism in cold-stored potato tubers with decreased expression of sucrose phosphate synthase. Plant, Cell & Environment. 1998;21(3):285–99.

64. Clasen BM, Stoddard TJ, Luo S, Demorest ZL, Li J, Cedrone F, et al. Improving cold storage and processing traits in potato through targeted gene knockout. Plant biotechnology journal. 2016;14(1):169–76.

65. Wang S, Zhang S, Wang W, Xiong X, Meng F, Cui X. Efficient targeted mutagenesis in potato by the CRISPR/Cas9 system. Plant Cell Reports. 2015;34(9):1473–6.

66. Butler NM, Baltes NJ, Voytas DF, Douches DS. Geminivirus-mediated genome editing in potato (Solanum tuberosum L.) using sequence-specific nucleases. Frontiers in plant science. 2016;7:1045.

67. Forsyth A, Weeks T, Richael C, Duan H. Transcription activator-like effector nucleases (TALEN)-mediated targeted DNA insertion in potato plants. Frontiers in plant science. 2016;7:1572.

68. Butler NM, Atkins PA, Voytas DF, Douches DS. Generation and inheritance of targeted mutations in potato (Solanum tuberosum L.) using the CRISPR/Cas system. PloS one. 2015;10(12):e0144591.

69. Andersson M, Turesson H, Nicolia A, Fält A-S, Samuelsson M, Hofvander P. Efficient targeted multiallelic mutagenesis in tetraploid potato (Solanum tuberosum) by transient CRISPR-Cas9 expression in protoplasts. Plant cell reports. 2017;36(1):117–28.

70. Schattenberg JM, Bergheim I. Nutritional intake and the risk for non-alcoholic fatty liver disease (NAFLD). Multidisciplinary Digital Publishing Institute; 2019.

71. Park J-YK, Koehler KM. Probabilistic Quantitative Assessment of Coronary Heart Disease Risk From Dietary Exposure to Industrially Produced Trans-Fatty Acids in Partially Hydrogenated Oils. Toxicological Sciences. 2019;172(1):213–24.

72. Haun W, Coffman A, Clasen BM, Demorest ZL, Lowy A, Ray E, et al. Improved soybean oil quality by targeted mutagenesis of the fatty acid desaturase 2 gene family. Plant biotechnology journal. 2014;12(7):934–40.

73. al Amin N, Ahmad N, Wu N, Pu X, Ma T, Du Y, et al. CRISPR-Cas9 mediated targeted disruption of FAD2-2 microsomal omega-6 desaturase in soybean (Glycine max. L). BMC biotechnology. 2019;19(1):9.

74. Zhang P, Du H, Wang J, Pu Y, Yang C, Yan R, et al. Multiplex CRISPR/Cas9-mediated metabolic engineering increases soya bean isoflavone content and resistance to soya bean mosaic virus. Plant biotechnology journal. 2019.

75. Li Z, Liu Z-B, Xing A, Moon BP, Koellhoffer JP, Huang L, et al. Cas9-guide RNA directed genome editing in soybean. Plant physiology. 2015;169(2):960–70.

76. Kanazashi Y, Hirose A, Takahashi I, Mikami M, Endo M, Hirose S, et al. Simultaneous site-directed mutagenesis of duplicated loci in soybean using a single guide RNA. Plant cell reports. 2018;37(3):553–63.

77. Ito Y, Nishizawa-Yokoi A, Endo M, Mikami M, Toki S. CRISPR/Cas9-mediated mutagenesis of the RIN locus that regulates tomato fruit ripening. Biochemical and Biophysical Research Communications. 2015;467(1):76–82.

78. Hu N, Xian Z, Li N, Liu Y, Huang W, Yan F, et al. Rapid and user-friendly open-source CRISPR/Cas9 system for single-or multi-site editing of tomato genome. Horticulture research. 2019;6(1):7.

79. Hare PD, Chua N-H. Excision of selectable marker genes from transgenic plants. Nature biotechnology. 2002;20(6):575.

80. Schaart JG, Krens FA, Pelgrom KT, Mendes O, Rouwendal GJ. Effective production of marker-free transgenic strawberry plants using inducible site-specific recombination and a bifunctional selectable marker gene. Plant biotechnology journal. 2004;2(3):233–40.

81. Corneille S, Lutz K, Svab Z, Maliga P. Efficient elimination of selectable marker genes from the plastid genome by the CRE-lox site-specific recombination system. The Plant Journal. 2001;27(2):171–8.

82. Sun N, Zhao H. Transcription activator-like effector nucleases (TALENs): A highly efficient and versatile tool for genome editing. Biotechnology and bioengineering. 2013;110(7):1811–21.

83. Ma C, Liu M, Li Q, Si J, Ren X, Song H. Efficient BoPDS gene editing in cabbage by the CRISPR/Cas9 system. Horticultural Plant Journal. 2019;5(4):164–9.

84. Zhang K, Nie L, Cheng Q, Yin Y, Chen K, Qi F, et al. Effective editing for lysophosphatidic acid acyltransferase 2/5 in allotetraploid rapeseed (Brassica napus L.) using CRISPR-Cas9 system. Biotechnology for biofuels. 2019;12(1):1–18.

85. Xu Z-S, Feng K, Xiong A-S. CRISPR/Cas9-mediated multiply targeted mutagenesis in orange and purple carrot plants. Molecular biotechnology. 2019;61(3):191–9.

86. Klimek-Chodacka M, Oleszkiewicz T, Baranski R. Visual Assay for Gene Editing Using a CRISPR/Cas9 System in Carrot Cells. Plant Genome Editing with CRISPR Systems: Springer; 2019. p. 203–15.

87. Chen K, Gao C. Developing CRISPR technology in major crop plants. Advances in New Technology for Targeted Modification of Plant Genomes: Springer; 2015. p. 145–59.

88. Belhaj K, Chaparro-Garcia A, Kamoun S, Nekrasov V. Plant genome editing made easy: targeted mutagenesis in model and crop plants using the CRISPR/Cas system. Plant methods. 2013;9(1):39.

89. Agapito-Tenfen SZ, Okoli AS, Bernstein MJ, Wikmark O-G, Myhr AI. Revisiting risk governance of GM plants: the need to consider new and emerging gene-editing techniques. Frontiers in plant science. 2018;9.

90. Bryksin AV, Matsumura I. Overlap extension PCR cloning: a simple and reliable way to create recombinant plasmids. Biotechniques. 2010;48(6):463–5.

91. Wang M, Liu Y, Zhang C, Liu J, Liu X, Wang L, et al. Gene editing by co-transformation of TALEN and chimeric RNA/DNA oligonucleotides on the rice OsEPSPS gene and the inheritance of mutations. PLoS One. 2015;10(4):e0122755.

92. Mahfouz MM, Piatek A, Stewart Jr CN. Genome engineering via TALENs and CRISPR/Cas9 systems: challenges and perspectives. Plant biotechnology journal. 2014;12(8):1006–14.

